## Supplementary Figures for "*N*^6^-methyladenosine modification is not a general trait of viral RNA genomes"

Baquero-Pérez *et al*.

### Supplementary Information

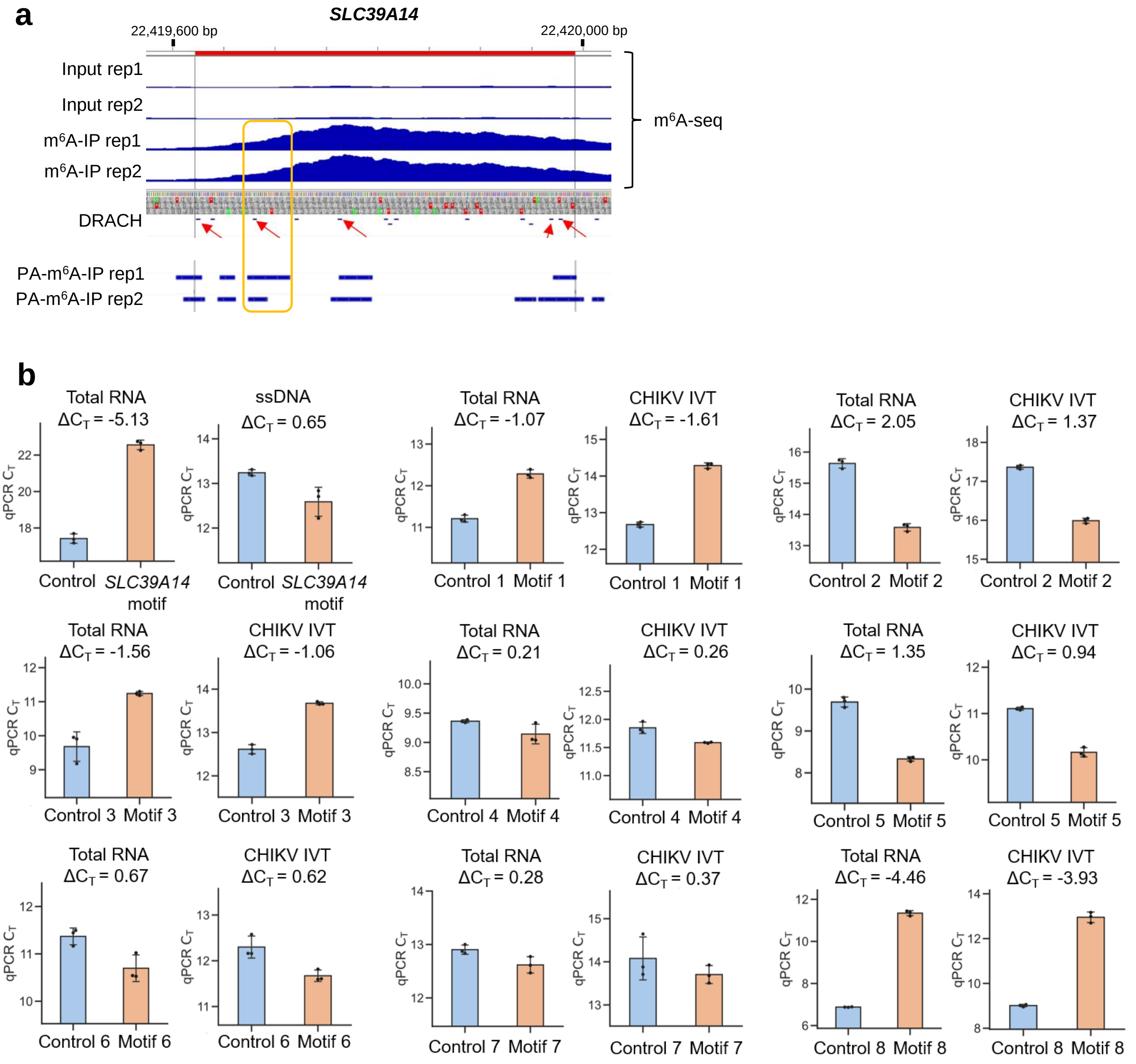


**Supplementary figure 1. Raw $C_T$ cycles from SELECT analyses of CHIKV RNAs. a)** Genome browser tracks display our m$^6$A-seq dataset along with publically available PA-m$^6$A-seq datasets[27]. Mapped reads for the *SLC39A14* transcript are shown. DRACH motifs are depicted as blue thin lines spanning the transcript. Red arrows indicate regions where DRACH motifs overlap with both biological replicates of PA-m$^6$A-seq. The positive control modified adenosine in the *SLC39A14* mRNA (genomic coordinate Chr8:22419678) is highlighted with a yellow line. **b)** The bar chart presents mean values obtained from three independent SELECT reactions, with the error bars representing the standard deviation (SD.). Results are representative from two independent infections.

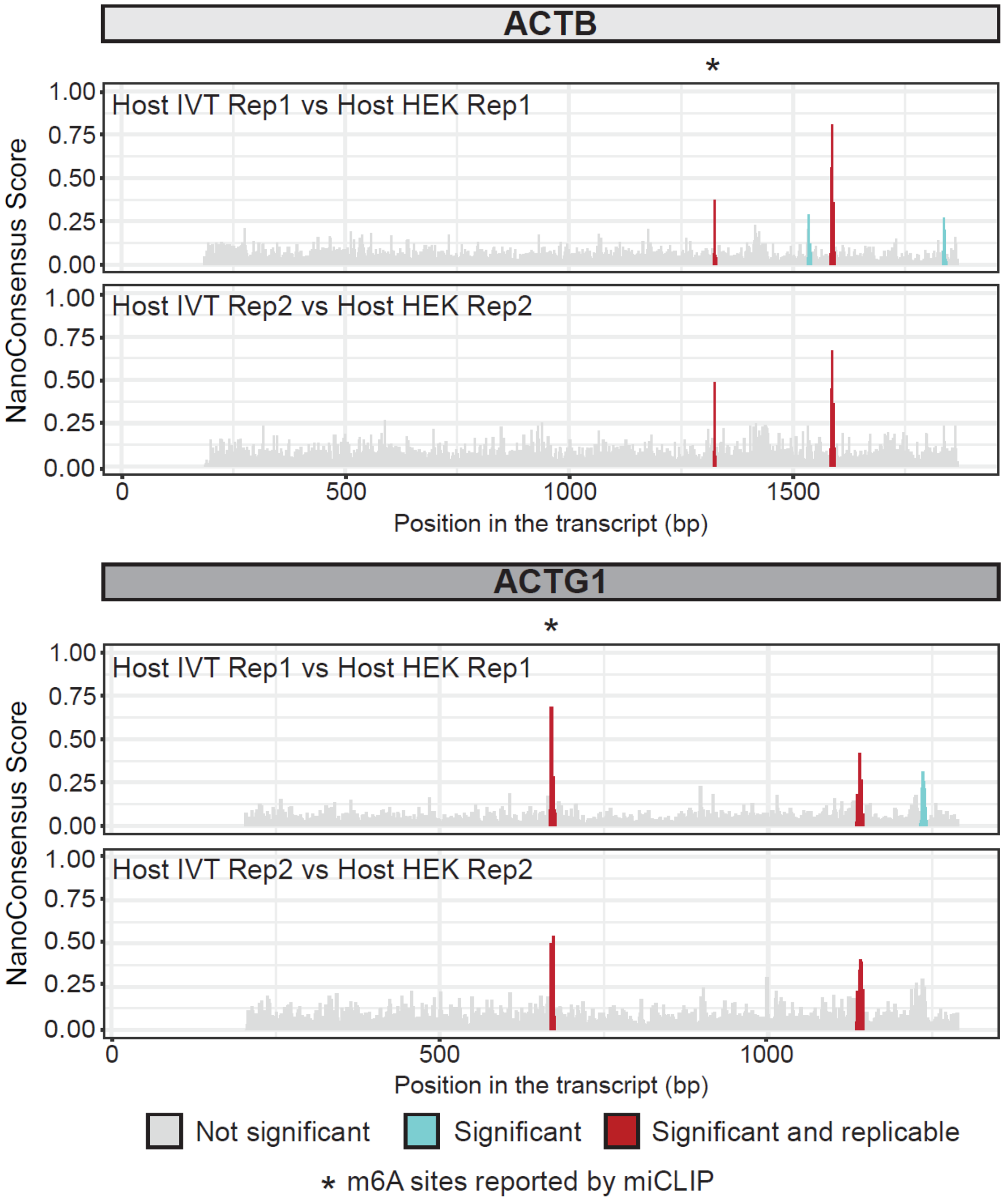


**Supplementary figure 2. DRS identifies differentially modified sites in host mRNAs.** *NanoConensus*' scores across the human transcripts *ACTB* (upper panels) and *ACTG1* (lower panels) when comparing reads from the CHIKV-infected samples and an IVT human transcriptome. *NanoConsensus´* default parameters were used. In grey, non-significant positions; in blue, regions identified by *NanoConsensus* in only one replicate; in red, regions identified in both replicates. $m^6A$ sites previously identified by miCLIP[28] are indicated with an asterisk (*).

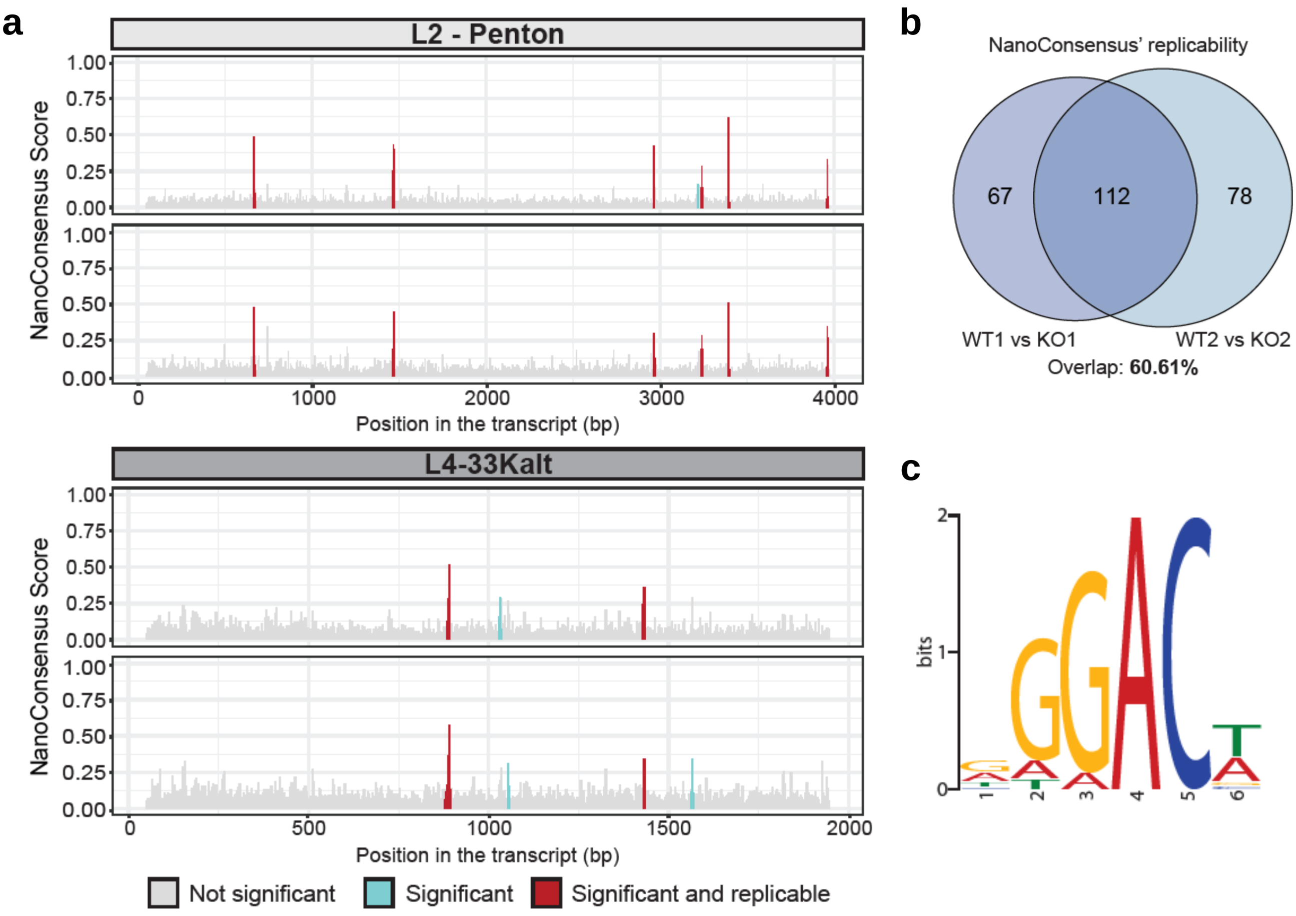


**Supplementary figure 3. *NanoConsensus* can identify modified sites in the** Ad5 **transcriptome. a)** *NanoConensus*' scores across adenovirus' transcripts L2-Penton (upper panels) and L4-33Kalt (lower panels) when comparing reads from wild-type and METTL3 knockout samples. *NanoConsensus´* default parameters were used. In grey, non-significant positions; in blue, regions identified by *NanoConsensus* in only one replicate; in red, regions identified in both replicates. **b)** Venn diagram showing the replicability of sites predicted by *NanoConsensus* across comparisons. **c)** Motif significantly enriched in replicable sites identified by *NanoConsensus* in the adenovirus' transcriptome.

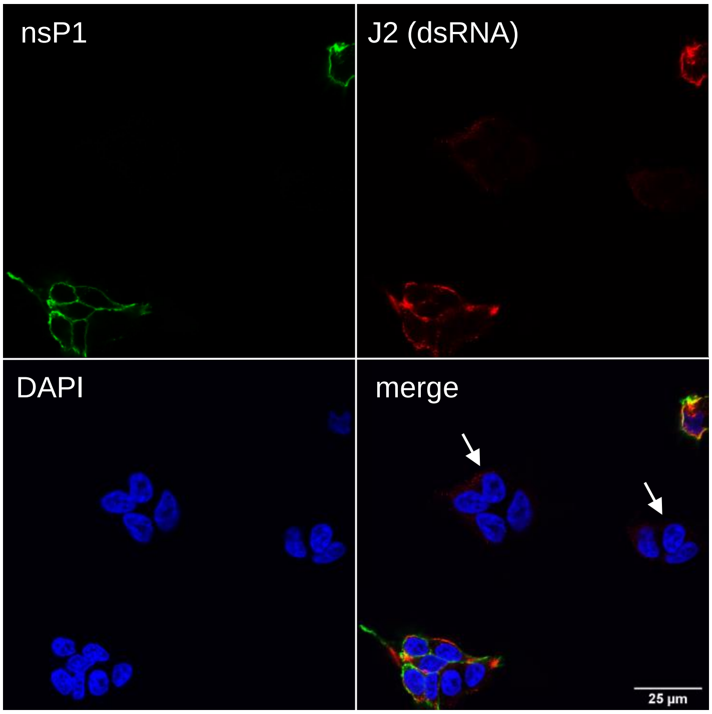


**Supplementary figure 4. J2 antibody recognises viral RNA in CHIKV-infected HEK293T cells.** Cells were infected for 18 hr at an MOI of 4. Infected cells were identified by labelling the CHIKV membrane-associated nonstructural protein 1 (nsP1). White arrows point to clusters of non-infected cells.

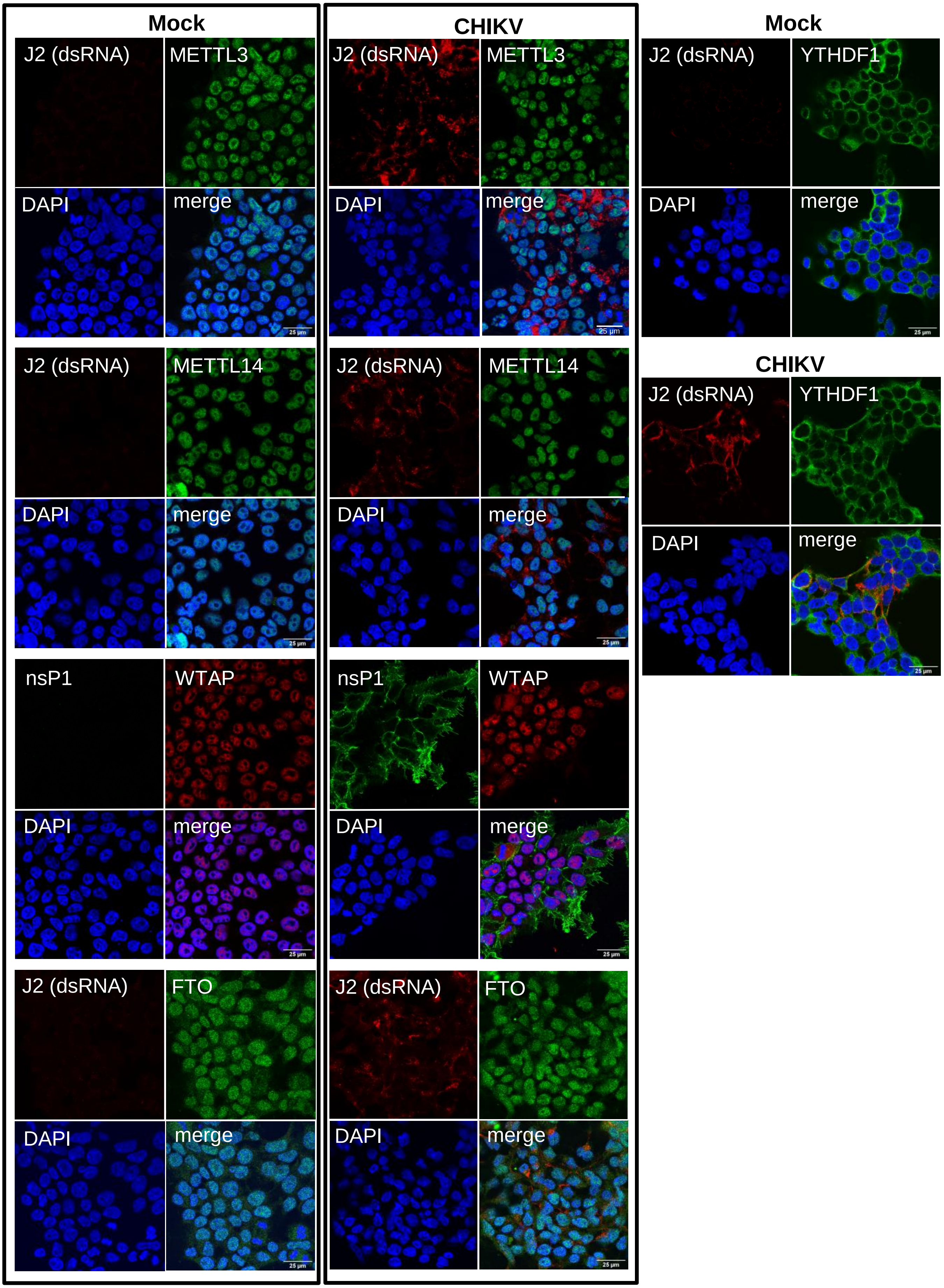


**Supplementary figure 5. Localization of METTL3, METTL14, WTAP, FTO, or YTHDF1 in mock- or CHIKV-infected HEK293T cells.** Cells were infected for 18 hr at an MOI of 4. Note that antibodies against METTL3, METTL14, FTO, YTHDF1, and nsP1 are derived from rabbits, while J2 and WTAP are derived from mice.

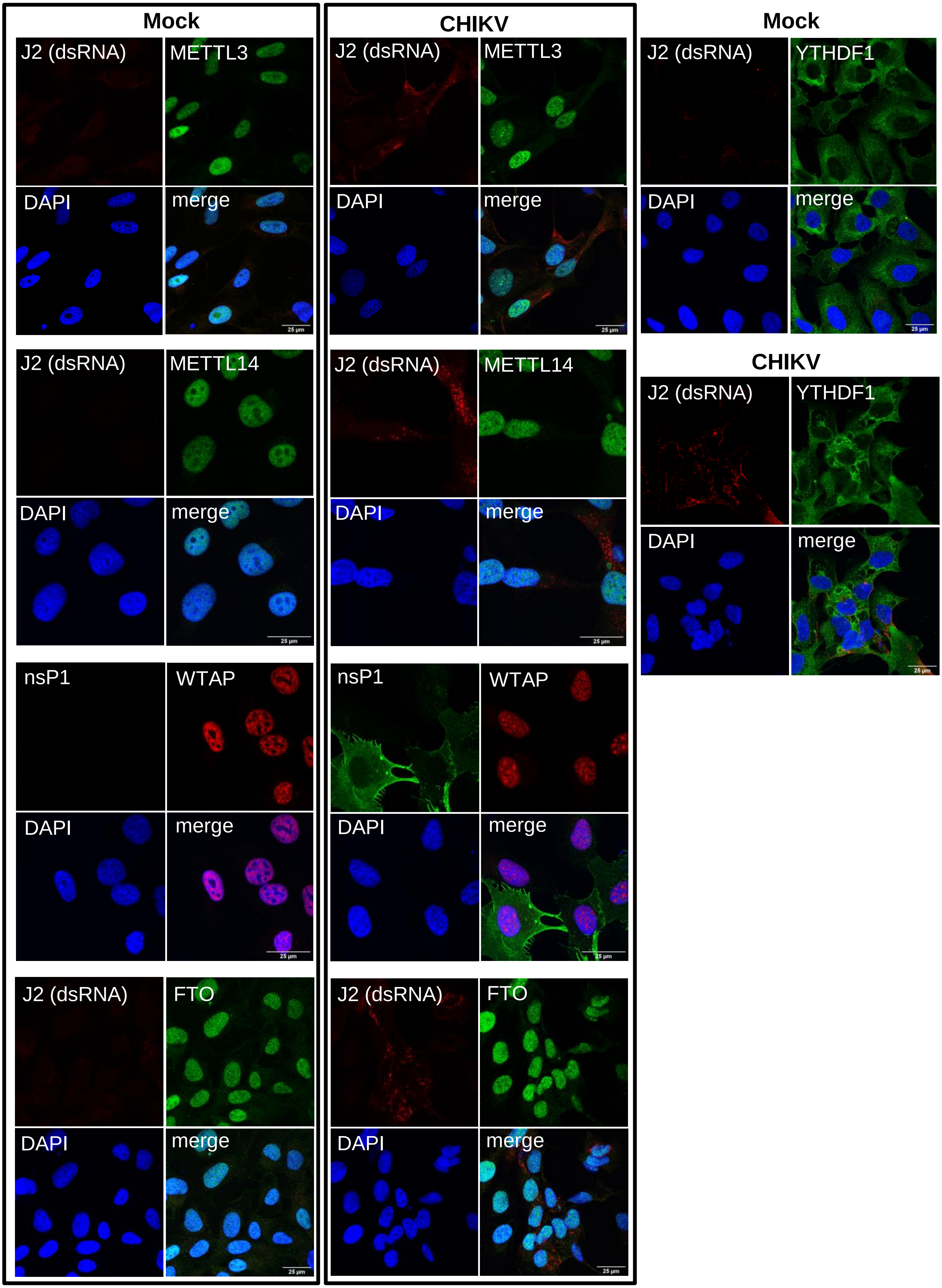


**Supplementary figure 6. Localization of METTL3, METTL14, WTAP, FTO, or YTHDF1 in mock- or CHIKV-infected U2OS cells.** Cells were infected for 18 hr at an MOI of 4. Note that antibodies against METTL3, METTL14, FTO, YTHDF1, and nsP1 are all derived from rabbits while J2 and WTAP are derived from mice.

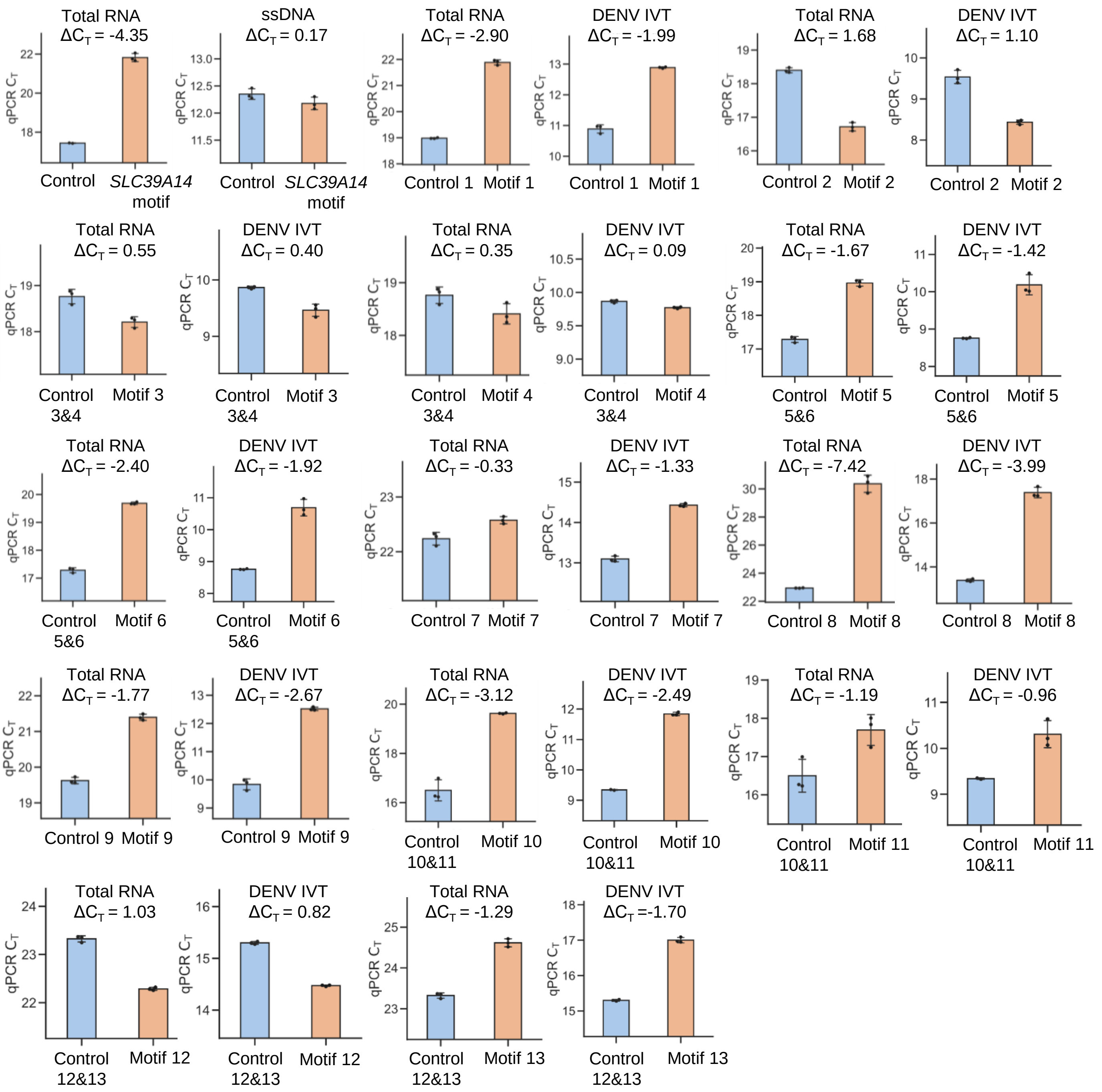


**Supplementary figure 7. Raw $C_T$ cycles from SELECT analyses of DENV RNAs in Huh7 cells.** The bar chart shows mean values from three SELECT reactions with the error bars showing SD.

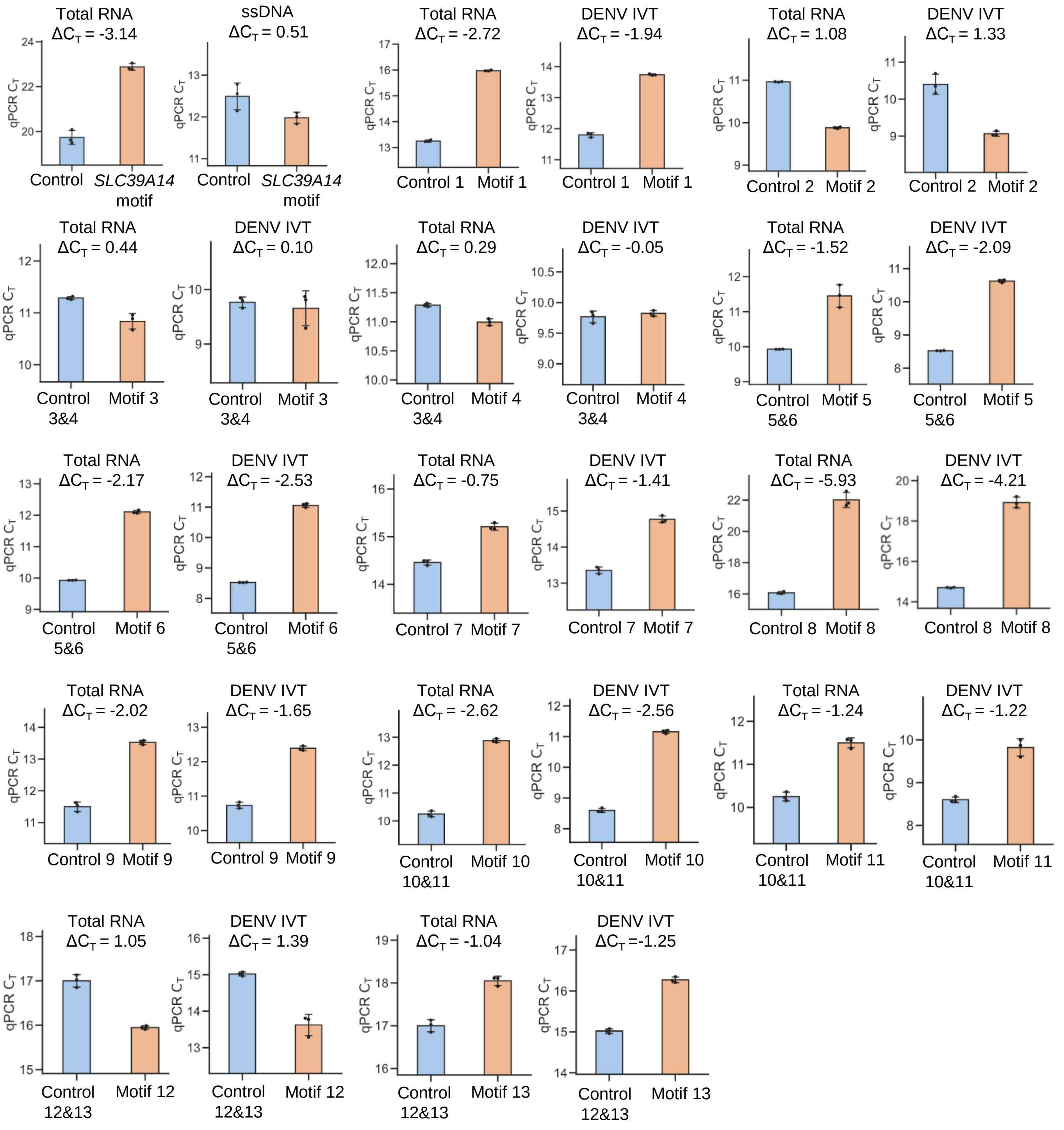


**Supplementary figure 8. Raw $C_T$ cycles from SELECT analyses of DENV RNAs in HEK293T cells.** The bar chart shows mean values from three SELECT reactions with the error bars showing SD.

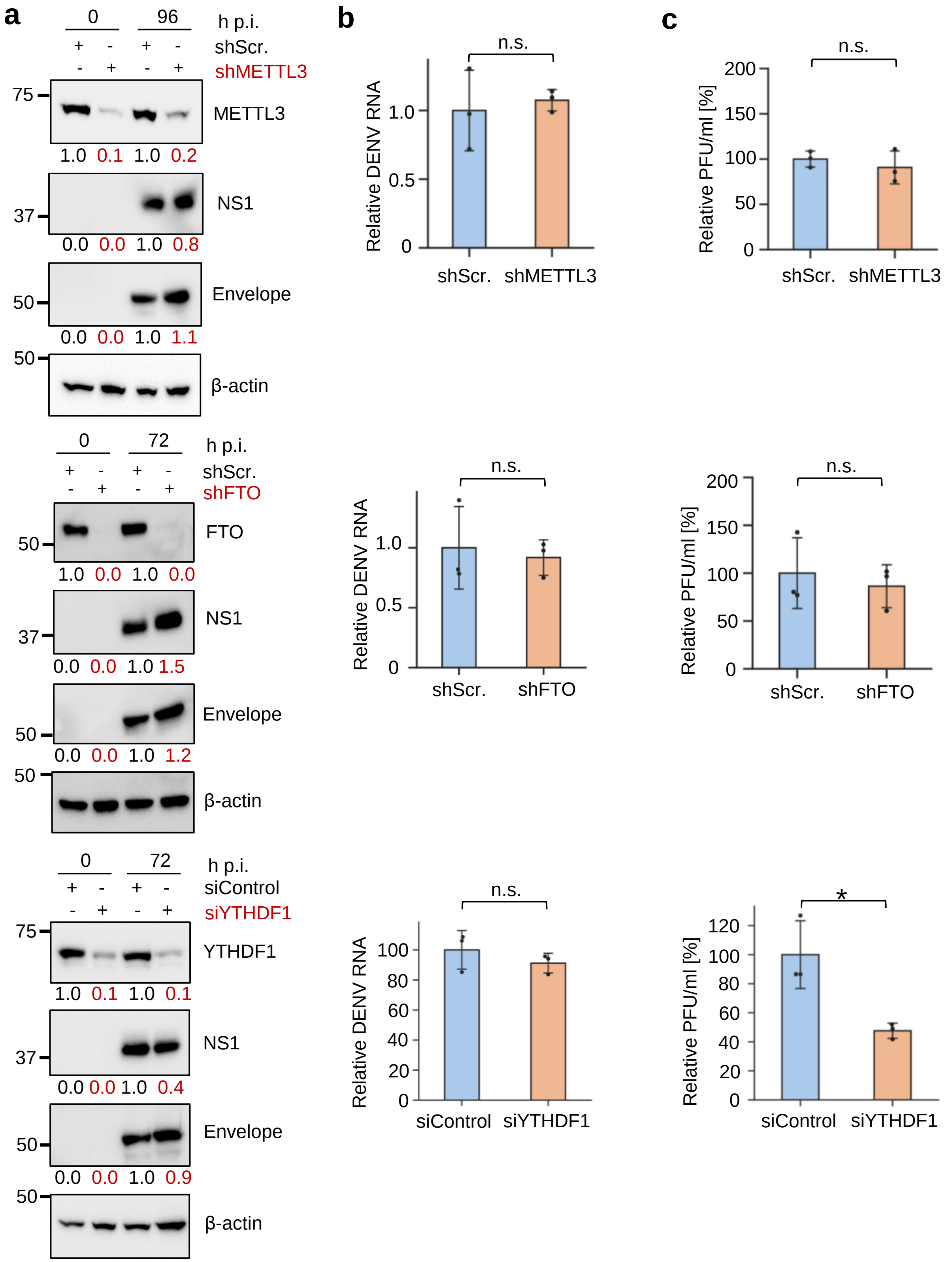


**Supplementary figure 9. Effect of depletion of METTL3, FTO, or YTHDF1 on DENV infection.** Stable shRNA HEK293T cell lines were infected at an MOI of 0.5 for the indicated time post-infection. At 24 hr of siRNA depletion, siRNA-treated HEK293T cells were infected for 72 hr at an MOI of 0.5. Scr. = scramble, h p.i. = hours post-infection. n.s. = not significant. **a)** Western blot quantification analyses are representative from 2 independent infections. β-actin-normalised values from depleted samples, below each band, are shown relative to their controls. **b)** Intracellular viral RNA levels were quantified by qRT-PCR and normalised against the housekeeping gene *GAPDH*. DENV RNA levels in depleted samples are shown relative to their corresponding control. The bar chart shows mean values from 3 independent infections with the error bars showing SD. **c)** Supernatants collected 72 h p.i. from DENV-infected control and knockdown cells were titered by plaque assay in HEK293T cells. The bar chart shows relative mean values from 3 independent replicates with the error bars showing SD. All statistical analyses were performed using a two-tailed *t*-test. n.s. = not significant.

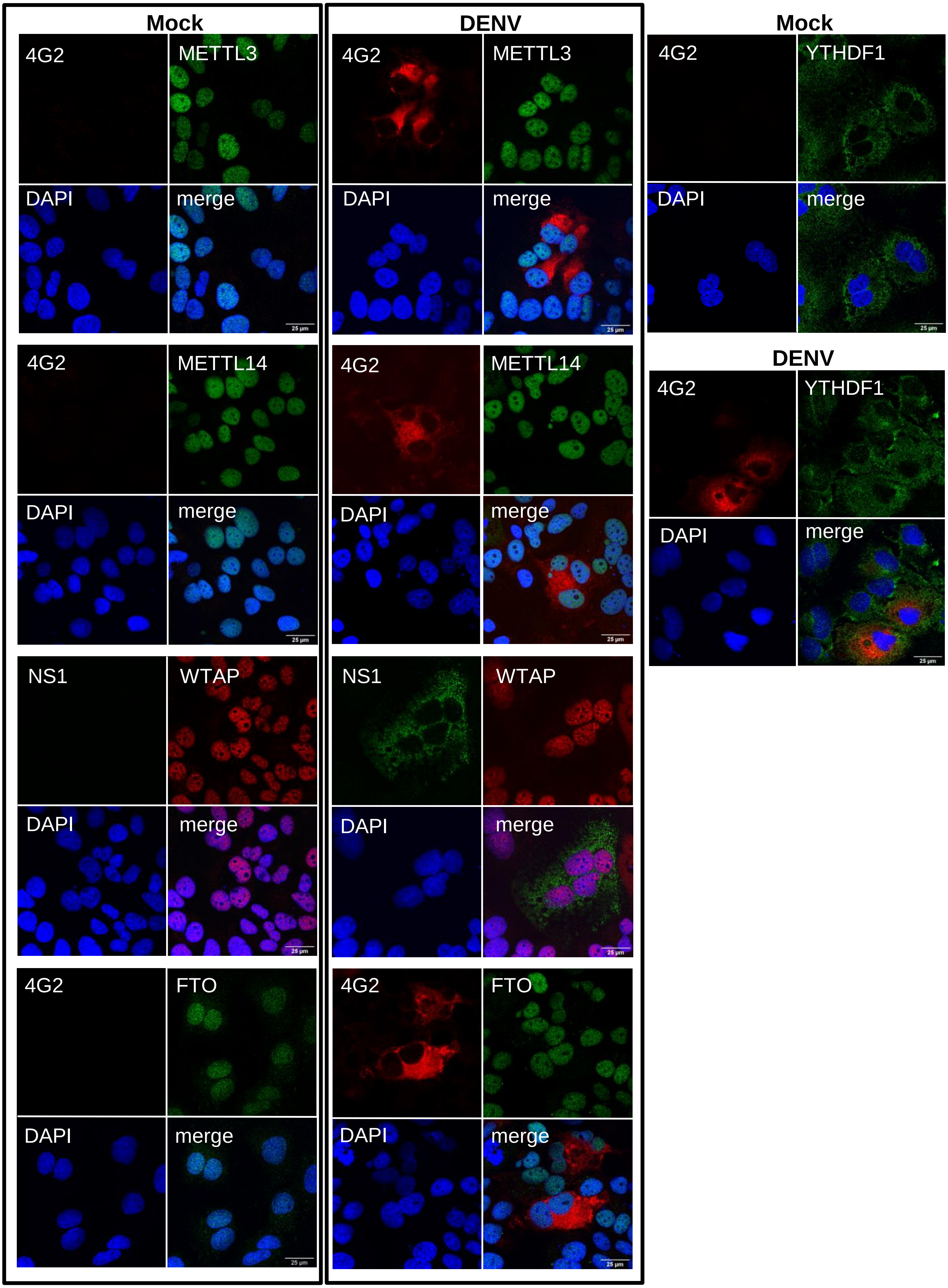


**Supplementary Fig. 10. Localization of METTL3, METTL14, WTAP, FTO, or YTHDF1 in mock- or DENV-infected Huh7 cells.** Cells were infected for 48 hr at an MOI of 3. 4G2 antibody recognises the DENV-2 envelope protein (E protein). NS1 antibody recognises the DENV-2 nonstructural protein 1 (NS1). Note that antibodies against METTL3, METTL14, FTO, YTHDF1, and NS1 are all derived from rabbits while WTAP and 4G2 are derived from mouse.
